# Amino acid-level differences in alpha tubulin sequences are uniquely required for meiosis

**DOI:** 10.1101/2024.10.11.617919

**Authors:** Li Chen, Xi Chen, Anna Kashina

**Affiliations:** Department of Biomedical Sciences, School of Veterinary Medicine, University of Pennsylvania, Philadelphia, PA 19104, USA; Department of Cell and Developmental Biology, Perelman School of Medicine, University of Pennsylvania, Philadelphia, PA 19104, USA

## Abstract

Tubulin is the major structural constituent of the microtubule cytoskeleton. Yeast *Schizosaccharomyces pombe* contain two α- tubulins genes, *nda2* and *atb2*, that are highly functionally distinct: *nda2* deletion is lethal, while lack of *atb2* does not interfere with cell viability. The functional determinants underlying this distinction are unknown. Here we used CRISPR-Cas9 gene editing to generate a yeast strain expressing Atb2 amino acid sequence utilizing Nda2 codon usage in the native Nda2 locus. Such Nda2-coded Atb2 (NCA) yeast, unlike Nda2 knockout, were viable and displayed no visible abnormalities in cell morphology or vegetative life cycle. However, these NCA yeast showed strong impairments in sporulation and meiosis, including major meiotic delays and high rates of abnormal chromosome segregation. Our data indicate that the amino acid sequence of Nda2 is uniquely required for normal meiosis, and identify a novel determinant that underlies functional distinction between closely related tubulin isoforms.

## INTRODUCTION

Tubulin is the major structural constituent of the microtubule cytoskeleton that plays an essential role in intracellular transport, cell division, and cellular homeostasis. In cells, obligatory α/ β- tubulin heterodimers serve as building blocks for microtubule assembly. Different eukaryotic species contain a variety of α- and β- tubulin genes that encode proteins with high sequence similarity. The mechanisms underlying such functional diversity of tubulin isoforms are unknown.

Earlier studies proposed that the existence of multiple genes encoding nearly identical proteins is a general safekeeping mechanism that ensures the functional robustness of highly essential proteins like tubulin through their duplication in the genome. These earlier studies predicted that tubulin isoforms are functionally redundant and can easily substitute for each other in case of disrupting mutations in one of the tubulin genes. However, more recent studies showed that tubulin isoforms have different properties [1, 2]. Furthermore, gene knockout studies in different organisms strongly suggest that tubulin isoforms play non-redundant functions in vivo. The underlying mechanisms driving this functional divergence, and the specific processes linked to unique features of tubulin isoforms, are unknown.

Studies of tubulin isoforms in mammals are highly complex, due to the combinatorial complexity of their composition in different tissues and physiological states. Important insights have been gained in budding yeast *Saccharomyces cerevisiae*, which encode a single beta tubulin and two highly homologous alpha tubulins that partly functionally overlap but also play non-redundant roles in such processes as spindle positioning [3] and regulation of microtubule dynamics [4]. However, studies of tubulin isoforms typically rely on gene deletions and don’t control for differences in the nucleotide sequence, which has recently come into focus as a potentially global determinant of cytoskeleton function [5]. Work from our lab recently demonstrated that nucleotide-level context and silent nucleotide substitutions play key roles in the global functions of actin isoforms, by regulating their translation dynamics, posttranslational modifications, and in vivo properties [6]. By extension, it appears likely that similar principles may apply to other protein isoform families, including tubulins. If so, controlling for the nucleotide-level variables appears to be an essential part of the studies of tubulin isoform functions. Yeast *Schizosaccharomyces pombe* represents a highly promising experimental model for addressing the functional diversity of tubulins while controlling for these variables. *S. pombe* is a well-established model organism that shares many physiological similarities with higher eukaryotes. The genome of *S. pombe* encodes one single β- tubulin and two α- tubulin genes (Fig. S1A), *nda2* and *atb2*, that are highly functionally distinct: *nda2* is essential for viability, while *atb2* deletion does not lead to any strong phenotypes. This limited complexity makes *S. pombe* an ideal organism to identify the determinants of differential tubulin function.

To address the nucleotide-versus amino acid-level differences driving the distinction between Nda2 and Atb2, we used CRISPR-Cas9 gene editing to generate a yeast strain expressing Nda2-coded Atb2 (NCA). In this strain, the Nda2 codon usage is preserved in its native locus, and is minimally edited to encode its close homolog Atb2 (Fig. S1B). Thus, any phenotypic changes in this strain would arise largely from the amino acid-level differences between these two proteins, enabling us to functionally separate the contribution of nucleotide sequence.

Strikingly, unlike *nda2* deletion, NCA yeast were viable and displayed no visible abnormalities in cell morphology, cell division, or cell growth. However, NCA cells showed strong impairments of meiosis, resulting in extremely poor ability to form proper spores. NCA meiotic spindles showed altered structure and dynamics, accompanied by reduced binding of the microtubule crosslinker Ase1, the spindle kinesins Klp2 and Klp9, suggesting that the differential ability of Nda2/Atb2 tubulins to bind these proteins modulates in meiosis. Our data indicate that the amino acid sequence of Nda2 is uniquely required for normal meiosis and identify a novel determinant that underlies functional distinction between closely related tubulin isoforms.

## RESULTS

### Replacement of Nda2 protein with Atb2 does not affect yeast viability or growth

To test the functional determinants driving differential functions of Nda2 and Atb2, we used CRISPR/Cas9 to edit the Nda2 gene to encode Atb2 protein, by introducing a series of minimal point mutations into the animo acid-coding codons while preserving the overall Nda2 codon usage (Fig. S1). In this yeast strain, termed NCA for “<u>N</u>da2-<u>C</u>oded-<u>A</u>tb2”, Nda2 gene is nearly intact, but all the Nda2 protein is completely absent, fully replaced with Atb2. This enabled us to address whether the nucleotide-dependent or the amino acid-dependent determinants of Nda2 – or both – are required for its essential cellular function.

Strikingly, in contrast to Nda2 gene deletion, NCA yeast were completely viable, despite lacking Nda2 protein. These yeast cells had normal morphology and growth and displayed no visible abnormalities in cell shape or size (Fig. 1). To test whether NCA yeast undergo normal division – a process that heavily depends on the microtubule-based mitotic spindle – we performed time lapse imaging of yeast cultures transformed with the fluorescent histone Hht2-GFP (the marker of the chromatin), Cut11-GFP (that labels the nuclear envelope) and Sid4-mCherry (the marker for the spindle pole bodies. Since *S. pombe* have closed mitosis and meiosis that occur without the nuclear envelope breakdown, these markers enable direct measurements of spindle length and observation of spindle dynamics (Fig 2).

**Figure 1.**
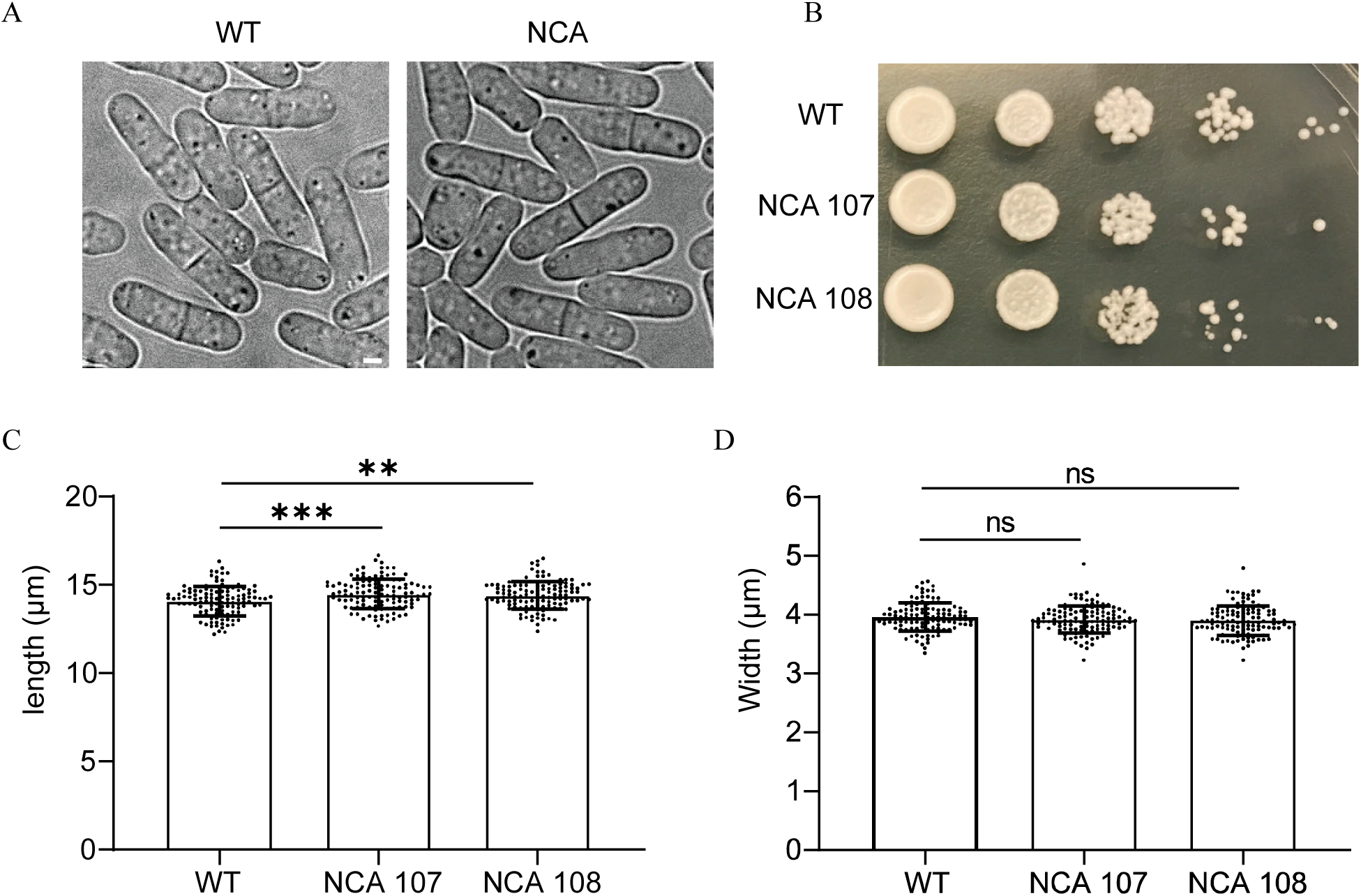
*Schizosaccharomyces pombe* cellular morphology and serial dilution assay between wild type (WT) and *nda2* coded *atb2* (NCA) strains: (A) Images show the similar cellular morphology between WT and NCA strains. (B) Serial dilution assay indicates no growth defects on NCA strains compared to WT. (C and D) Quantitative assay of cellular length and width (mean±SD) show the significant difference of cellular length between WT and NCA strains (WT, 14.09 ±0.83 μm, n=114; NCA 107, 14.51±0.83 μm, n=119; NCA 108, 14.42±0.78 μm, n=118), however no difference of cellular width are detected between WT and NCA (WT, 3.96 ±0.24 μm, n=114; NCA 107, 3.92±0.23 μm, n=119; NCA 108, 3.9±0.25 μm, n=118). (Scale bar, 2 μm), (** P<0.01, *** P<0.001).

**Figure 2.**
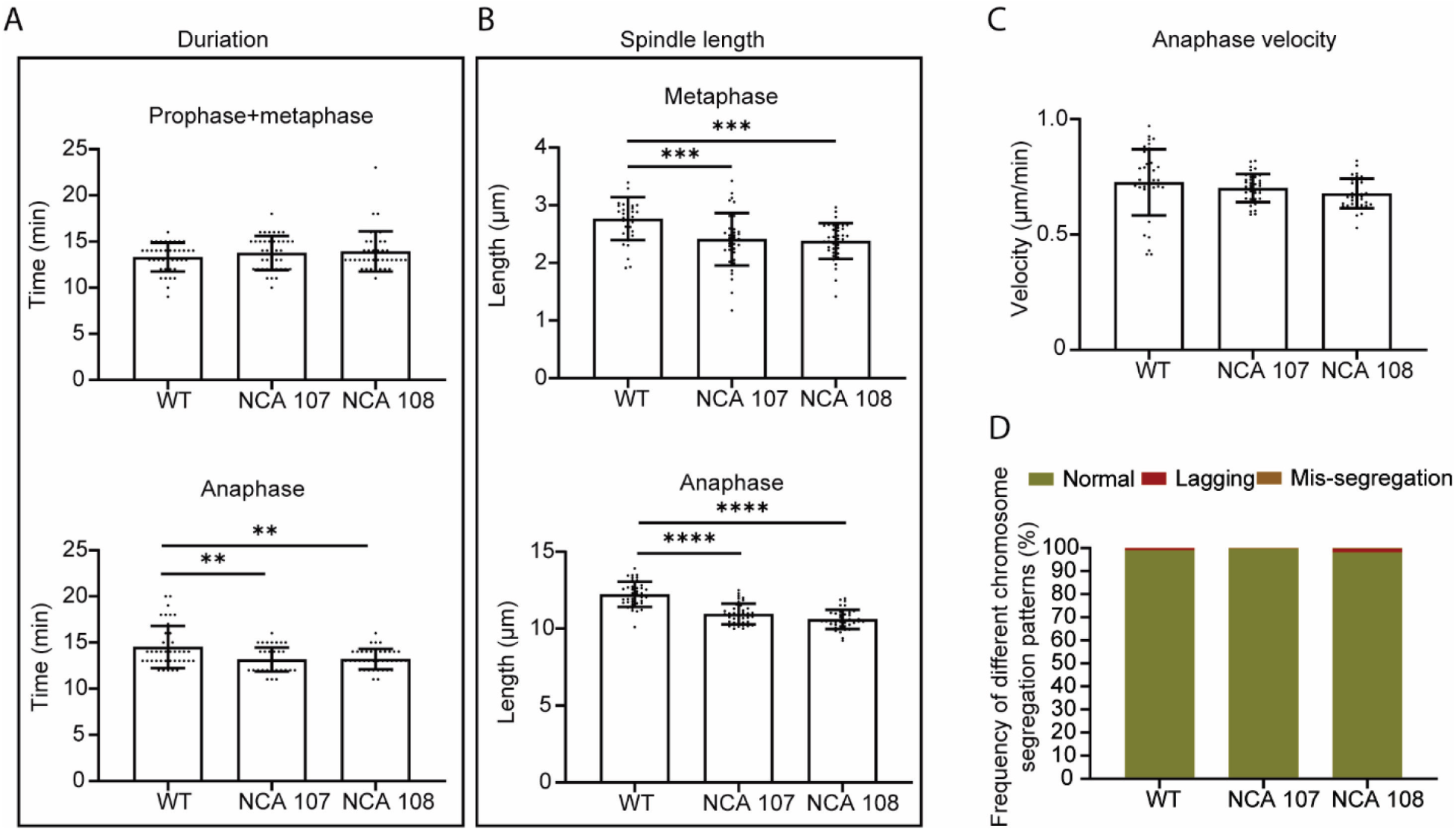
Cellular mitosis analyses between WT and NCA strains: (A) Duration of different phase in mitosis. Prophase to metaphase: WT, 13±2 min, n=40; NCA 107, 14±2 min, n=40; NCA 108, 14±2 min, n=39. Anaphase: WT, 15±2 min, n=40; NCA 107, 13±1 min, n=40; NCA 108, 13±1 min, n=39. (B) Mitosis spindle length in different phase. Metaphase: WT, 2.77±0.37 µm, n=40; NCA 107, 2.41±0.46 µm, n=40; NCA 108, 2.39±0.3 µm, n=39. End of anaphase: WT, 12.23±0.8 µm, n=40; NCA 107, 10.96±0.67 µm, n=40; NCA 108, 10.61±0.63 µm, n=39. (C) Anaphase velocity analysis shows no difference between WT and NCA strains: WT, 0.73±0.14 µm/min, n=40; NCA 107, 0.7±0.06 µm/min, n=40; NCA 108, 0.68±0.06 µm/min, n=39. (D) Comparative plot of frequency of chromosome segregation defects in WT and NCA strains shows normal chromosome segregation in NCA strains (WT, n=203; NCA 107, n=203; NCA 108, n=218).

Measurements of different mitotic stages in actively dividing cells revealed no strong differences in the duration of pro-, meta-, or anaphase, and no substantial differences in the metaphase or anaphase spindle length or anaphase velocity. A statistically significant reduction of anaphase duration and spindle length was observed in both NCA strains, however this difference was very small, making it difficult to evaluate its potential biological effects. No difference in chromosome segregation was observed between NCA and control. Overall, despite the minor statistical differences in our measurements, we observed no substantial mitotic defects in NCA strains.

Thus, replacement of Nda2 protein with Atb2 does not lead to changes in yeast viability and vegetative life cycle, suggesting that these functions are fully dependent on Nda2’s nucleotide, rather than amino acid sequence.

### Replacement of Nda2 protein with Atb2 leads to strong impairments in sporulation

In addition to the vegetative cycle, microtubules play an essential role during meiosis, which is required for yeast sporulation in response to stress conditions in the environment. During sporulation, yeast undergo two subsequent meiotic divisions (meiosis I and meiosis II) to form four spores from each diploid yeast cell. Abnormalities in meiosis lead to abnormal spore count, with one, two, or three spores formed from each cell. Mitotic and meiotic spindle are believed to function in very similar ways, however a number of important differences exist between these two processes [7]. Thus, it is possible that tubulin isoform replacement in yeast may specifically alter meiosis – and consequently, sporulation -- without affecting any other microtubule-dependent processes. To test this, we cultured NCA and control yeast in the sporulation media and counted the frequency of zygotes with abnormal spore count in each culture.

Over 94% of wild type yeast formed normal zygotes with four spores each. In contrast, two independently obtained NCA yeast strains (NCA107 and NCA108) both showed very high frequency of abnormal sporulation, with approximately 50% of all zygotes containing less than four spores (Fig. 3). Thus, lack of Nda2 protein in these cells leads to highly prominent sporulation defects.

**Figure 3.**
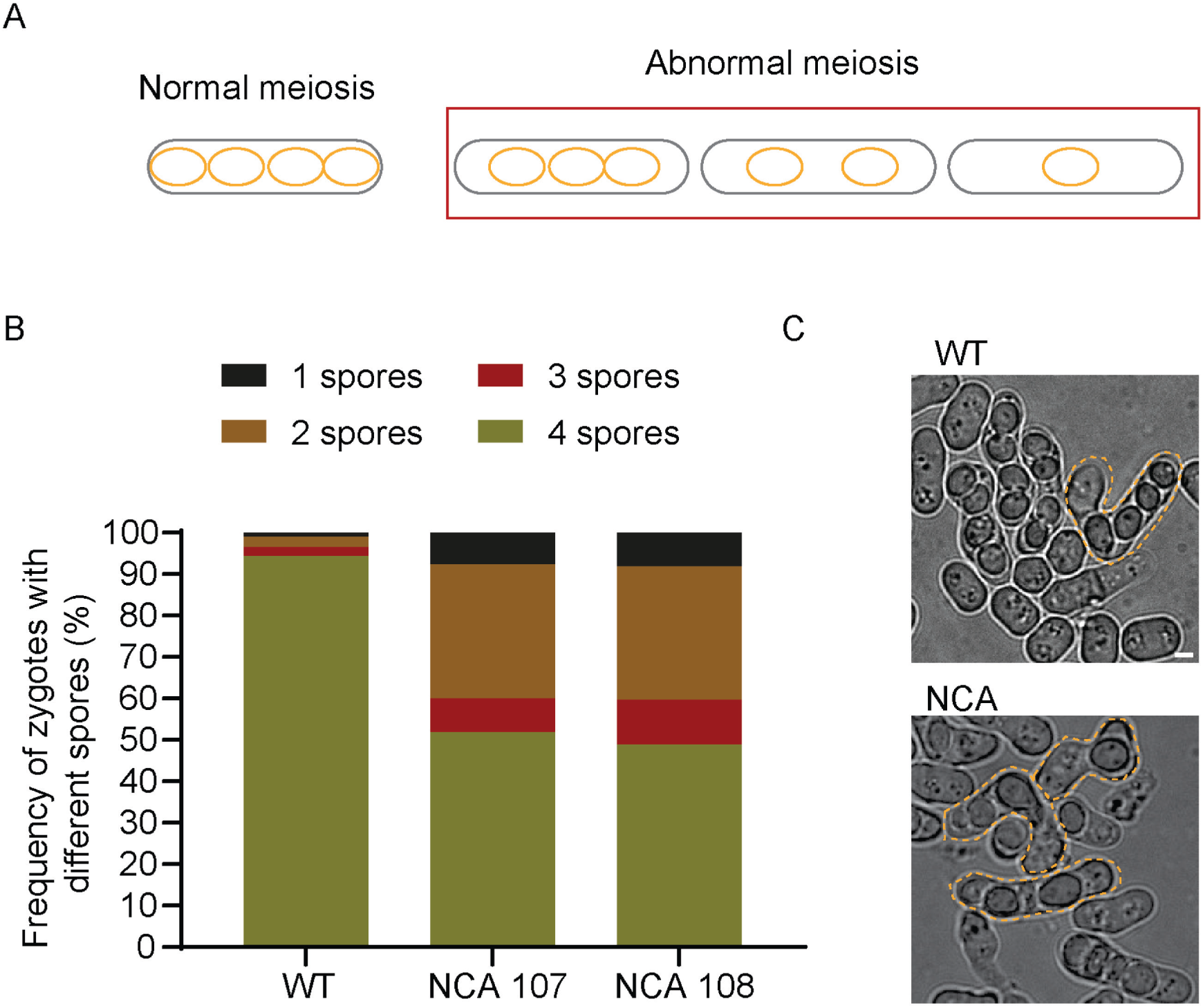
Sporulation assay of WT and NCA strains. (A) Illustration demonstrates the normal and abnormal spore formation of the sporulation process. (B) Comparison of frequency of abnormal sporulation between WT and NCA strains show large population of abnormal spores in cell from NCA strains (WT, abnormal sporulation 6%, n=1194; NCA 107, abnormal sporulation 52%, n=786; NCA 108, abnormal sporulation 49%, n=626;). (C) images of sporulated cells from WT and NCA strains show the normal and abnormal sporulation, yellow dashed lines outline the shape of cells. (Scale bar, 2µm).

### NCA yeast exhibit defects in meiotic chromosome segregation and spindle dynamics

To test directly whether meiotic chromosome segregation is abnormal in NCA cells, we performed time lapse imaging of meiosis in yeast cultures transformed with the fluorescently labeled histone Hht2-GFP (Fig. 4C). Strikingly, this imaging revealed substantial delays in both meiosis I and meiosis II, accompanied by frequent defects in chromosome positioning and movement, including abnormal chromosome lagging, mis-segregation, or failed segregation (Fig. 4 A, B).

**Figure 4.**
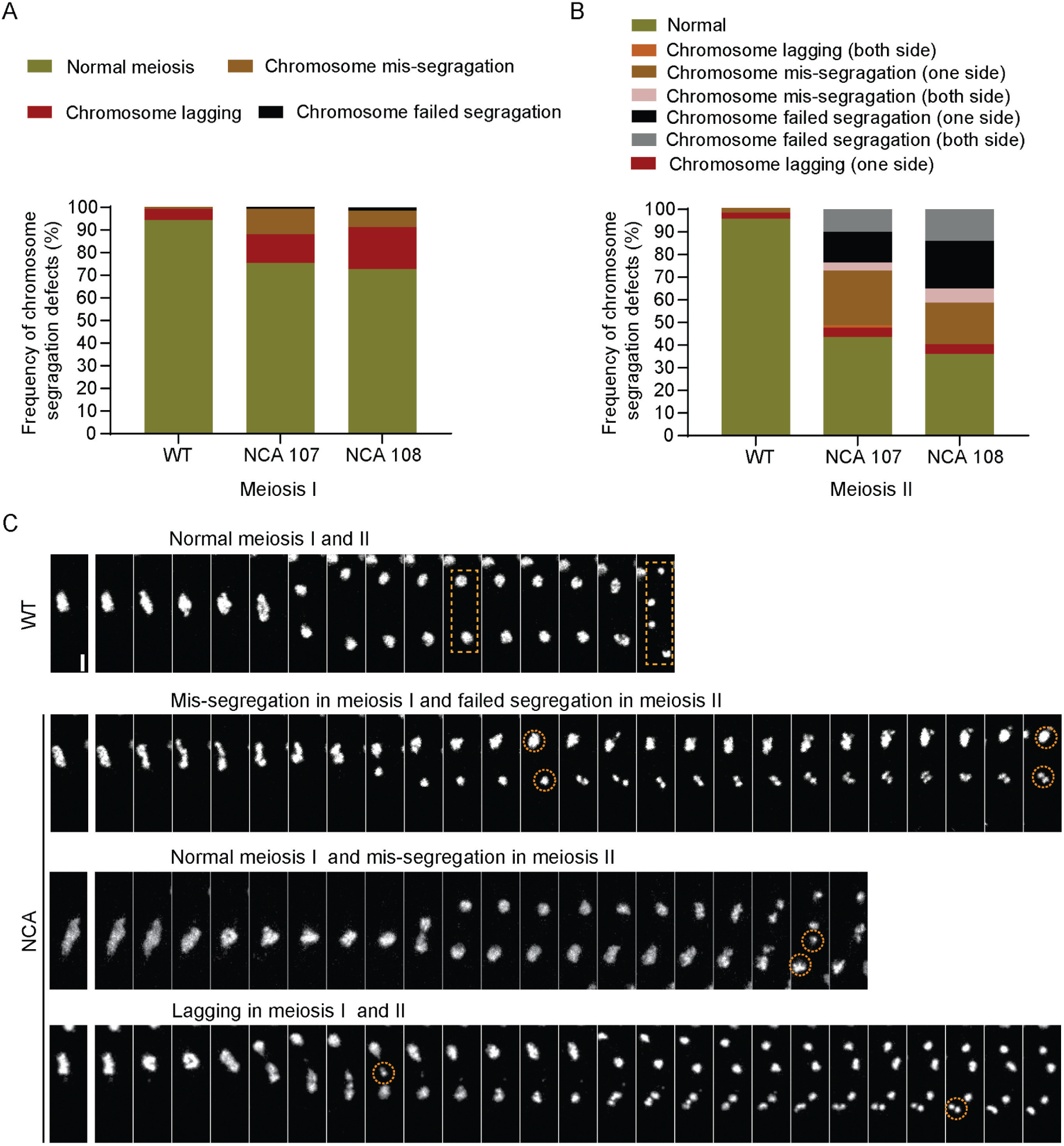
Chromosome segregation analyses through meiosis from WT and NCA strains. (A) meiosis I analysis shows that impaired chromosome segregation in NCA strains compared to WT (WT, 6% of total abnormal chromosome segregation, n=193; NCA 107, 25% of total abnormal chromosome segregation, n=174; NCA 108, 27% of total abnormal chromosome segregation, n=135). (B) column graph exhibits large portion of chromosome segregation in NCA strains during meiosis II (WT, 4% of total abnormal chromosome segregation, n=195; NCA 107, 47% of total abnormal chromosome segregation, n=170; NCA 108, 64% of total abnormal chromosome segregation, n=114). (C) Montage of normal chromosome segregation in WT and various of abnormal chromosome segregation in NCA strains expressing Hht2-GFP. Dashed rectangles indicate the normal chromosomes through the meiosis I and II, dashed cycles highlight the abnormal chromosomes during the meiosis I and II. (Scale bar, 2µm; maximum projection image of 7 focal planes with1-µm spacing; time interval of the slices is 6 min).

We next analyzed the morphology and dynamic of the meiotic spindles using live imaging of cells transformed with Cut11-GFP and Sid4-mCherry, the markers of nuclear envelope and spindle pole bodies, respectively (Fig. 5A). This imaging provided a more detailed picture of the meiosis dynamics. The duration of metaphase in both meiosis I and meiosis II were over two times higher in NCA yeast (Fig. 5B), indicating a substantial metaphase delay. In addition, we observed a dramatic shortening of meiotic spindle length (Fig. 5C), and over two-fold reduction in anaphase elongation velocity (Fig. 5D). The sum of these defects accounted for the fact that anaphase in NCA yeast took similar amount of time to control, since chromosomes in NCA yeast traveled approximately half the distance at half the speed. Plotting the meiotic spindle length change over time revealed substantial differences in overall spindle dynamics, with visible delays at all stages of meiosis I and II in NCA yeast (Fig. 5E). Overall, NCA yeast took roughly 1.5 times longer to complete meiosis: ∼60 minutes for meiosis I and ∼40 minutes for meiosis II, compared to ∼44 minutes for meiosis I, and ∼30 minutes for meiosis II seen in wild type control.

**Figure 5.**
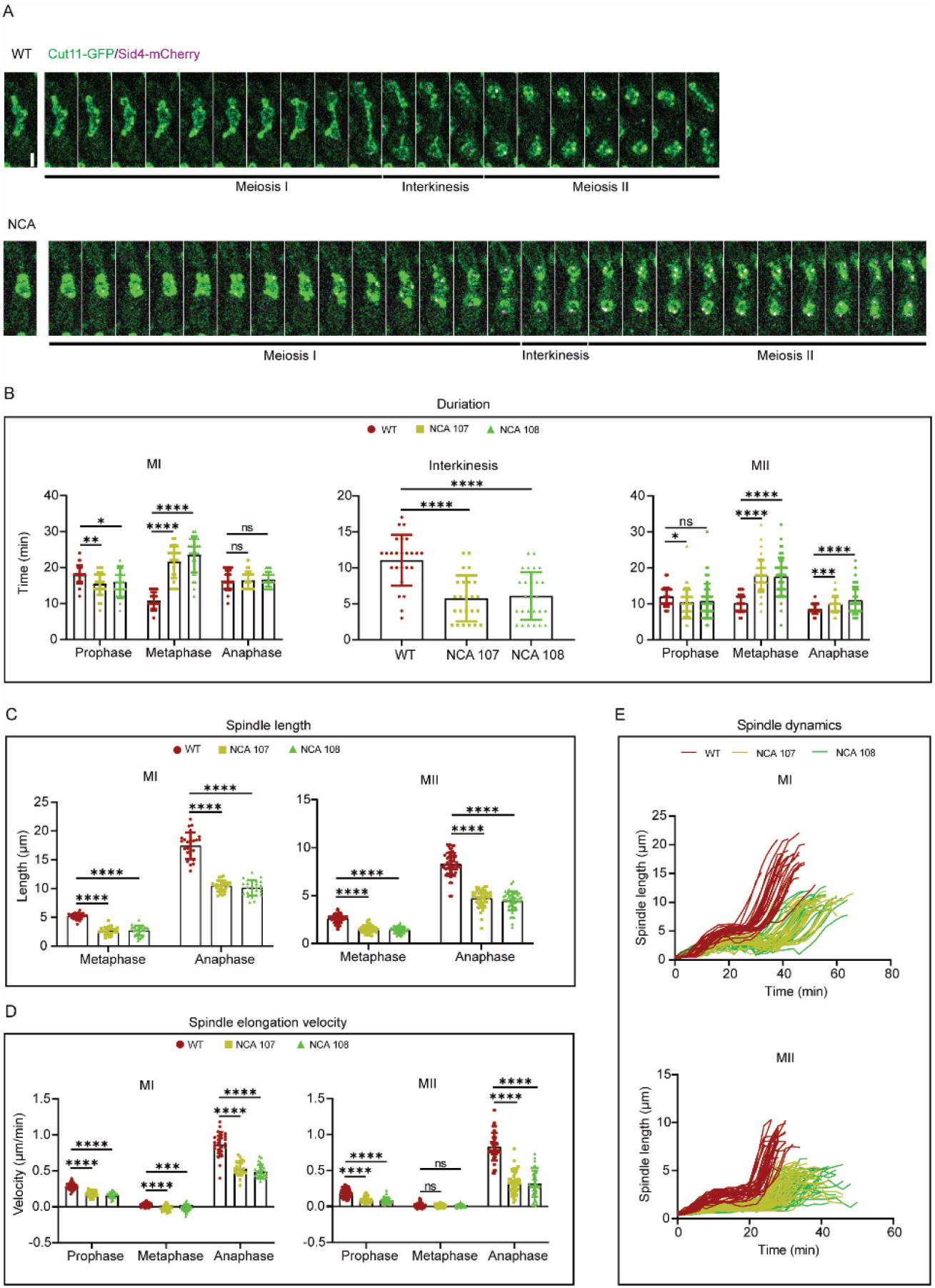
Characterization of spindle dynamics during meiosis I and II (MI and MII) in WT and NCA strains. (A) Time-lapse images of a meiotic spindle in cells expressing Cut11-GFP and Sid4-mCherry from WT and NCA strains. (B) Different phase duration time of meiosis. Meiosis I, prophase: WT, 18±3 min, n=25; NCA 107, 16±3 min, n=24; NCA 108, 16±4 min, n=26; metaphase: WT, 10±2 min, n=25; NCA 107, 21±5 min, n=24; NCA 108, 23±5 min, n=26; anaphase: WT, 16±3 min, n=25; NCA 107, 17±2 min, n=24; NCA 108, 17±2 min, n=26. Meiosis II, prophase: WT, 12±3 min, n=60; NCA 107, 10±4 min, n=43; NCA 108, 11±6 min, n=50; metaphase: WT, 10±2 min, n=60; NCA 107, 18±4 min, n=43; NCA 108, 17±5 min, n=50; anaphase: WT, 8±1 min, n=60; NCA 107, 10±2 min, n=43; NCA 108, 11±4 min, n=50. (C) Spindle length in meiosis. Meiosis I, metaphase: WT, 5.21±0.52 µm, n=25; NCA 107, 2.65±0.68 µm, n=24; NCA 108, 2.71±0.84 µm, n=26; end of anaphase: WT, 17.43±2.31 µm, n=25; NCA 107, 10.45±0.95 µm, n=24; NCA 108, 10.13±1.33 µm, n=26. Meiosis II, metaphase: WT, 2.61±0.45 µm, n=60; NCA 107, 1.52±0.39 µm, n=43; NCA 108, 1.45±0.33 µm, n=50; end of anaphase: WT, 8.27±1.18 µm, n=60; NCA 107, 4.69±0.91 µm, n=41; NCA 108, 4.46±0.97 µm, n=49. (D) Spindle elongation velocity of meiosis. Meiosis I, prophase: WT, 0.28±0.05 µm/min, n=25; NCA 107, 0.19±0.04 µm/min, n=24; NCA 108, 0.16±0.04 µm/min, n=24; metaphase: WT, 0.03±0.03 µm/min, n=25; NCA 107, -0.02±0.04 µm/min, n=24; NCA 108, -0.01±0.05 µm/min, n=26; anaphase: WT, 0.87±0.17 µm/min, n=25; NCA 107, 0.53±0.1 µm/min, n=24; NCA 108, 0.49±0.09 µm/min, n=26. Meiosis II, prophase: WT, 0.19±0.06 µm/min, n=60; NCA 107, 0.1±0.04 µm/min, n=43; NCA 108, 0.09±0.04 µm/min, n=50; metaphase: WT, 0.02±0.03 µm/min, n=60; NCA 107, 0.01±0.02 µm/min, n=43; NCA 108, 0.01±0.02 µm/min, n=50; anaphase: WT, 0.83±0.19 µm/min, n=60; NCA 107, 0.33±0.16 µm/min, n=40; NCA 108, 0.32±0.18 µm/min, n=49. (E) Comparative plots of spindle length versus time of WT and NCA strains in meiosis I and II. (Scale bar, 2 µm; maximum projection image of 7 focal planes with1-µm spacing; ns, non-significant; * P<0.05, ** P<0.01, **** P<0.0001; time interval between the slices is 4 min).

### NCA yeast exhibit defects in nuclear horsetail movement

Meiosis in fission yeast is preceded by a complex process of nuclear orientation, known as horsetail movement, when the elongated nucleus exhibits a series of back-and-forth oscillations required for chromosome positioning during the subsequent meiosis I. To test whether, in addition to the meiosis defects, NCA yeast also exhibited changes in the nuclear horsetail movement, we selected the time lapse videos of the events occurring 40 frames (80 minutes) earlier than the onset of meiosis I and tracked the movement of spindle pole bodies, marked with Sid4-mCherry, to record the range and velocity of nuclear movement (Fig. 6). Wild type spindle pole bodies during that time interval exhibited broad oscillating movements ranging around ∼4 um in length that gradually reduced toward the onset of meiosis (Fig 6A, red curve). In contrast, the moving distance of NCA spindle pole bodies ranged around 1.5 um (Fig. 6A, green curves), an over two-fold reduction compared to control (Fig. 6B, C). Visually, the nuclei at this stage exhibited a vastly different morphology: in wild type, the nuclei were elongated and clearly polarized, while the nuclei in NCA yeast looked more round and compact (Fig. 6D). This change likely reflects the fact that the NCA nuclei were not subjected to the same pushing and pulling forces as the wild type [8].

**Figure 6.**
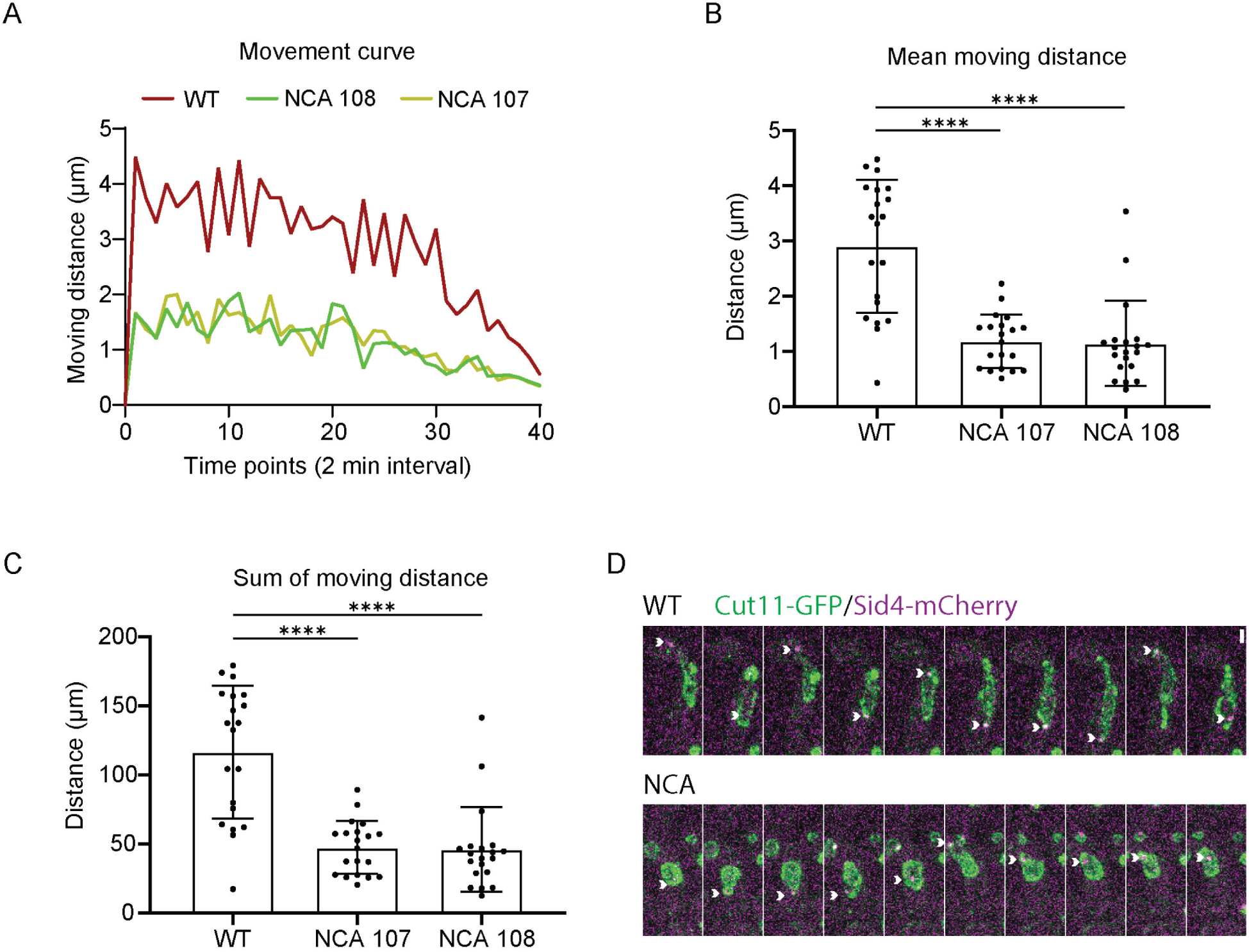
Nucleus horsetail movement analysis. (A) Comparative plots of moving distance versus time of WT and NCA strains, the time range for analysis was selected 40 time points before the onset of spindle pole bodies (PSB) separation. (B and C) Column graphs show the mean distance and sum distance of nucleus moving between WT and NCA strains (WT, mean 2.91±1.21 µm, sum 116.4±48.2 µm, n=20; NCA 107, mean 1.19±0.48 µm, sum 47.57±19.34 µm, n=20; NCA 108, mean 1.15±0.77 µm, sum 46.13±30.63 µm, n=20;). (D) Time-lapse images of nucleus horsetail movement in cells expressing Cut11-GFP and Sid4-mCherry from WT and NCA strains (white arrow heads indicate the SPB movement). (Scale bar 2 µm; maximum projection image of 7 focal planes with1-µm spacing; **** P<0.0001; time interval between the slices is 8 min).

### Meiotic spindles in NCA yeast have altered levels of key spindle proteins

To test the underlying molecular changes that may lead to the observed defects in meiotic chromosome segregation and spindle elongation, we analyzed the dynamic distribution of key proteins that are known to play a role in these processes throughout meiosis. We first focused on Ase1, a microtubule associated protein required for the bipolar spindle maintenance [9] that interacts with antiparallel microtubules to generate local scaffolds for the recruitment of key proteins involved in cell division [10]. We used live imaging of dividing yeast cells expressing Ase1-GFP and tracked its localization and distribution in wild type in NCA mutants. In wild type, Ase1-GFP localized to the spindle midzone in metaphase, consistent with prior observations and with its demonstrated role in maintaining spindle stability (Fig. 7A top row). In NCA, the Ase1-enriched spindle midzone was markedly shorter (Fig 7A bottom row), and the Ase1-GFP signal appeared paler than in control. Quantification of Ase1-GFP fluorescence levels revealed that in NCA yeast spindles contained ∼2 fold lower levels of Ase1 compared to wild type, both overall (Fig. 7C) as well as in meiosis I and II (Fig. 7D, E).

**Figure 7.**
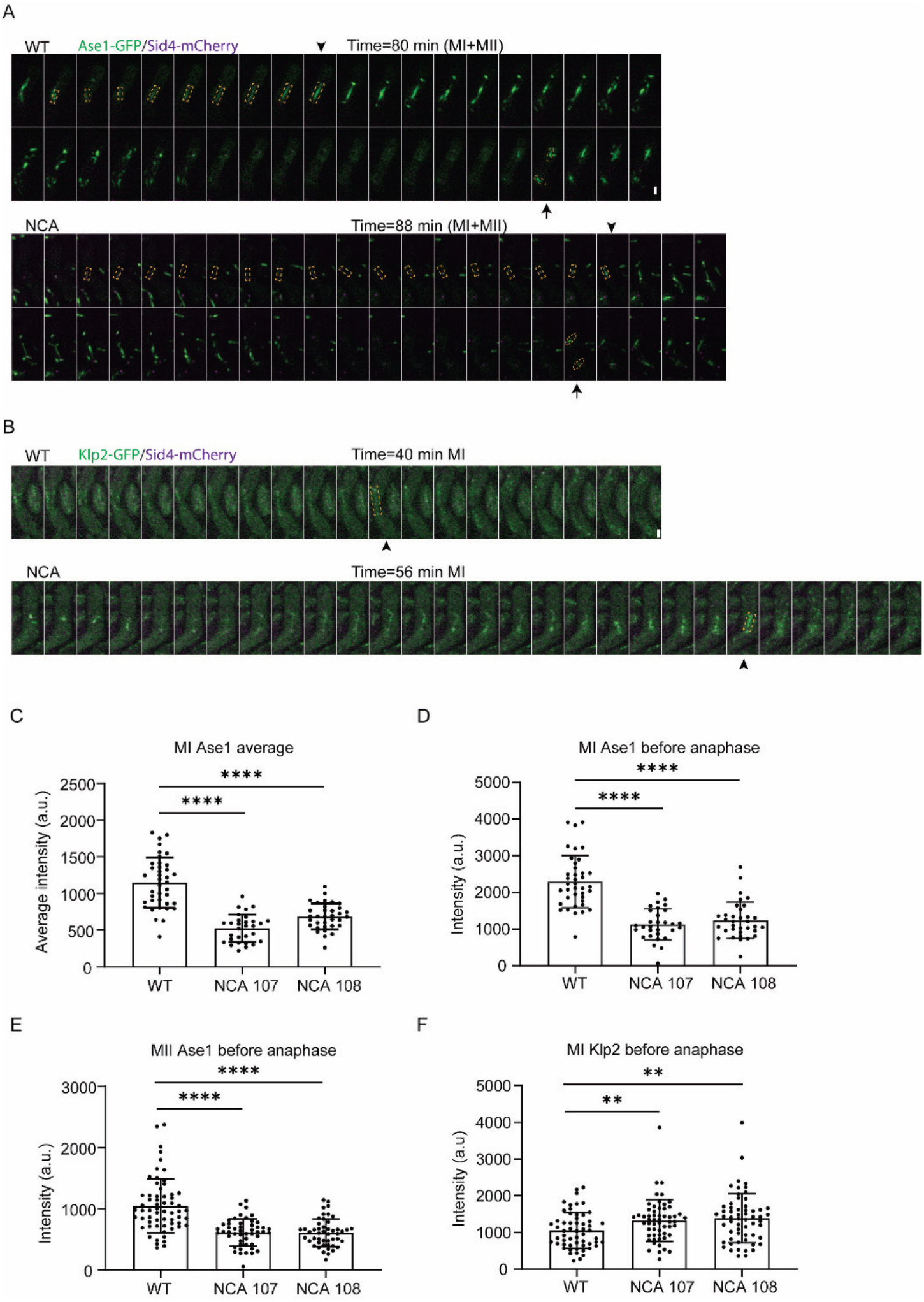
Quantitative analyses of Ase1 and Klp2 during meiosis I and II. (A) Time-lapse of WT and NCA cells expressing Ase1-GFP, dashed rectangles indicate the meiosis I Ase1-GFP intensity, arrow head indicates the time point that right before anaphase onset. Dashed ovals and arrow show the time point of Ase1-GFP intensity that right before anaphase onset during meiosis II. (B) Time-lapse of WT and NCA cells expressing Klp2-GFP, dashed rectangles and arrow head indicate the time point that right before anaphase onset in meiosis. (C) Average intensity of Ase1-GFP through prophase to metaphase in meiosis I (highlighted by dashed rectangles): WT, 1147±343.9, n=39; NCA 107, 526±187.7, n=30; NCA 108, 686.9±176.1, n=34. (D) Intensity of Ase1-GFP at the time point that right before anaphase onset (pointed by arrow head) in meiosis I: WT, 2296±713.9, n=39; NCA 107, 1130±424.7, n=30; NCA 108, 1245±493.7, n=34. (E) Intensity of Ase1-GFP at the time point that right before anaphase onset (pointed by arrow) in meiosis II: WT, 1051±438.6, n=62; NCA 107, 615.8±221.4, n=49; NCA 108, 609.2±224.8, n=48. (F) Intensity of Klp2-GFP at the time point that right before anaphase onset (pointed by arrow head) in meiosis I: WT, 1056±487.7, n=57; NCA 107, 1326±568.4, n=57; NCA 108, 1388±671, n=59 (Scale bar, 2 µm; sum projection image of 7 focal planes with1-µm spacing; ** P<0.01, **** P<0.0001; time interval between the slices is 4 min).

Ase1 acts as a key scaffold that recruits proteins essential for the spindle maintenance and progression of cell division in response to cyclin-dependent kinase/phosphatase cycles [11]. Some of its important partners include Klp2, a member of the kinesin 14 family that mediates microtubule bundling and sorting during spindle formation [12], and Klp9, a kinesin 6 family motor that mediates microtubule sliding and is recruited to the spindle by Ase1-mediated scaffolding [13]. Thus, Klp9 levels in the spindle directly depend on Ase1 interaction with the midzone microtubules, while Klp2 acts in concert but is not directly recruited by Ase1.

Measurements of the Klp2-GFP signal showed a minor but statistically significant increase in NCA yeast compared to control (Fig 7B, 7F). In contrast, Klp9-GFP levels showed an over 2-fold reduction in the spindle midzone of NCA yeast (Fig. 8). This reduction was proportional to the reduced Ase1 signal and persisted in both meiosis I (Fig. 8B, D) and meiosis II (Fig 8BC, E).

**Figure 8.**
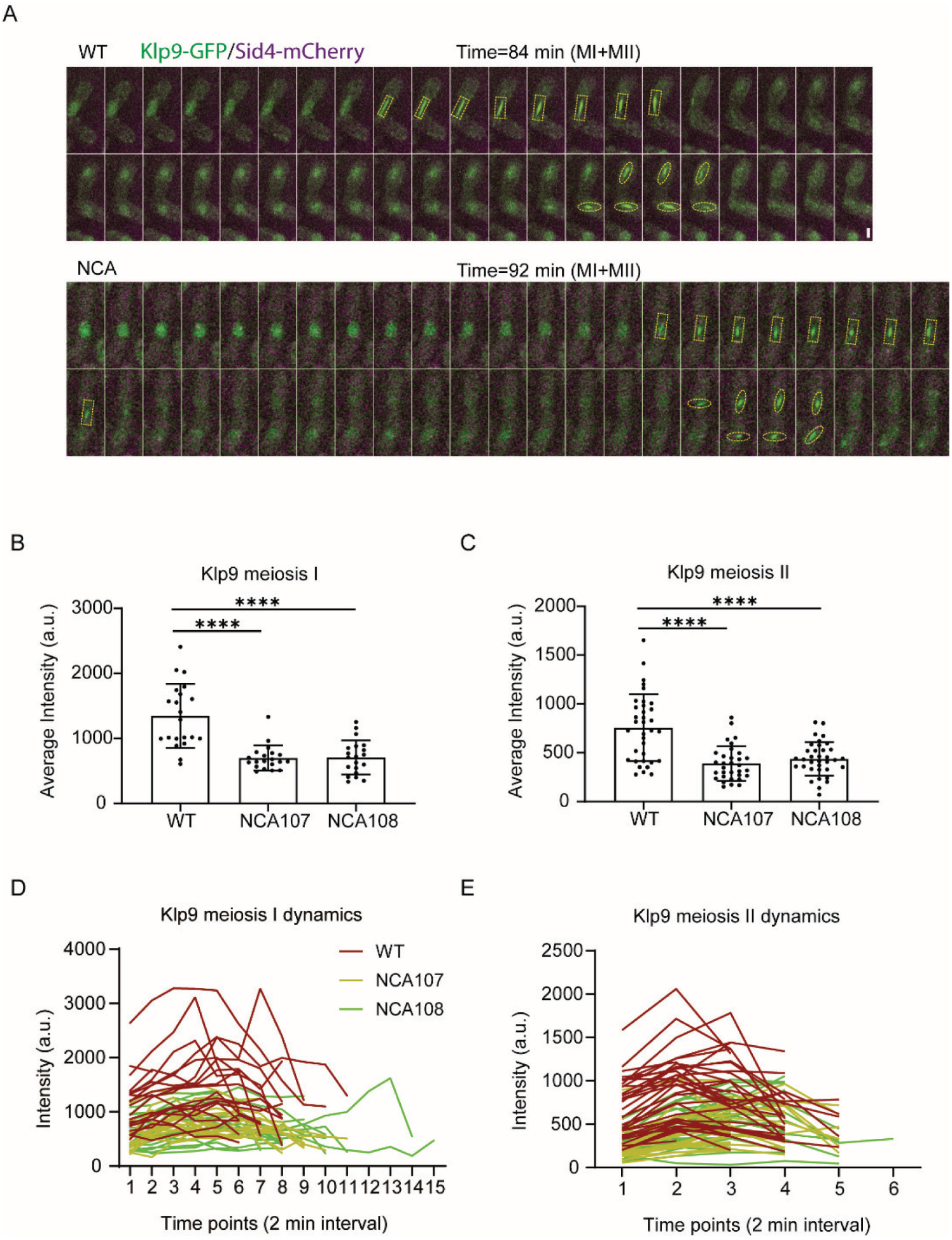
Quantitative analyses of Klp9 during meiosis I and II. (A) Time-lapse montages of WT and NCA cells expressing Klp9-GFP and Sid4-mCherry, dashed rectangles indicate the meiosis I Klp9-GFP intensity, dashed ovals show the Klp9-GFP intensity during meiosis II. (B and C) Comparative column graphs of Klp9-GFP average intensity in meiosis I and II. Meiosis I: WT, 1343±490, n=21; NCA 107, 698.5±192, n=20; NCA 108, 707.5±262.5, n=20. Meiosis II: WT, 756.4±342.5, n=35; NCA 107, 389.8±177.4, n=32; NCA 108, 437.1±171.7, n=35. (D and E) Comparative plot of Klp9-GFP intensity over time in meiosis I and II. (Scale bar, 2 µm; sum projection image of 7 focal planes with1-µm spacing; **** P<0.0001; time interval between the slices is 4 min).

Thus, NCA yeast show changes in key proteins required for the normal function of the meiotic spindle.

### NCA yeast exhibit changes in microtubule dynamics

Another prominent class of proteins required for the normal spindle function are microtubule end-binding proteins, including Mal3, the yeast homolog of the microtubule plus end tracking protein EB1 and a common marker for microtubule dynamics. To test whether lack of Nda2 protein in yeast leads to changes in microtubule dynamics, we performed live imaging of microtubules in yeast cells transfected with Mal3-GFP. First, we measured the overall microtubule growth rate, defined as the velocity of movement of the Mal3-decorated microtubule plus ends. This rate was dramatically reduced in NCA cells, indicating a strong defect in microtubule dynamics (Fig. 9A).

**Figure 9.**
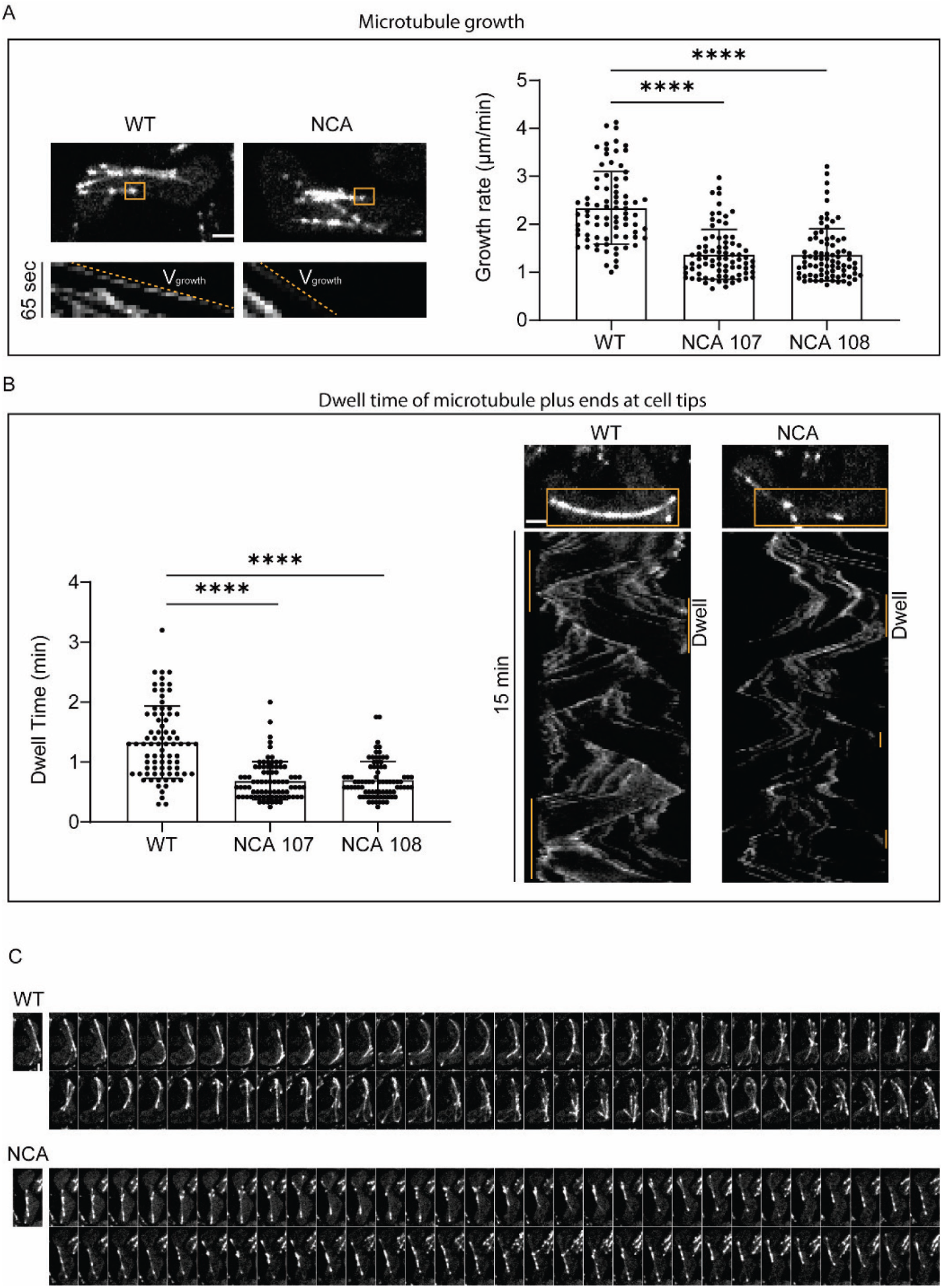
Interphase microtubule dynamics before meiosis onset. (A) Microtubule (MT) growth. Left: kymograph of MT growth over times in WT and NCA cells expressing Mal3-GFP. Top: yellow-box indicate the MT for growth monitoring. Bottom: kymograph generated from the yellow-boxed area; dashed lines indicate the MT growth trajectory. Right: MT growth rate in WT and NCA strains: WT, 2.34±0.76 µm/min, n=80; NCA 107, 1.37±0.52 µm/min, n=80; NCA 107, 1.37±0.54 µm/min, n=80. (B) Dwell time of MT plus ends at cell tips. Left: Kymograph of MT dwell time in WT and NCA cells expressing Mal3-GFP. Top: yellow-box indicate the area for kymograph generation. Bottom: kymograph generated from the yellow-boxed area; vertical lines indicate the MT dwell time. Right: MT tip dwell time from WT and NCA strains: WT, 1.3±0.6 min, n=80; NCA 107, 0.7±0.3 min, n=80; NCA 108, 0.7±0.3 min, n=80 (C) Time-lapse montages demonstrate the different MT behaviors between WT and NCA strains. (Scale bar, 2 µm; maximum projection image of 3 focal planes with1-µm spacing; time interval between the slices is 15 sec)

Next, we imaged the microtubule behavior at the cell tips during the time interval preceding the meiosis onset, when the microtubule plus ends dwell at the cell tips, enabling the generation of the major pushing force for nuclear horsetail movement [8]. The length of the dwell time is expected to be directly proportional to the pushing force, and to the extent of the nuclear movement. Quantification of this dwell time from the kymographs of the Mal3-GFP signal showed it to be significantly reduced in NCA yeast compared to control (Fig. 9B). By comparing to the WT cells, the NCA cells display shorter and non-curved microtubules indicate impaired microtubule stability (Fig. 9C). This observation is consistent with the reduced range and velocity of nuclear horsetail movement we observed in NCA yeast (Fig. 6).

## DISCUSSION

Our study for the first time separates the nucleotide- and amino acid-dependent functions of Nda2, the essential yeast alpha tubulin, and identifies its unique amino acid-level role in mediating normal meiosis. Deletion of Nda2 gene is lethal to yeast, but editing this gene to express another tubulin protein, Atb2, does not affect yeast viability or growth. At the same time, lack of Nda2 proteins in these yeast cells specifically affects meiotic division, resulting in meiotic delays and abnormal chromosome segregation that eventually lead to high rates of abnormal sporulation. These effects are linked to changes in microtubule dynamics and reduced binding of essential microtubule-associated proteins to the meiotic spindle. The present findings expand our understanding of the nucleotide- and amino acid-driven tubulin functions and outline a previously unknown mechanistic role for tubulin that specifically targets meiosis.

Many of the previous studies of the roles of tubulin isoforms relied on genetic knockouts or gene replacement, without controlling for amino acid versus nucleotide sequence. Using actin isoform family as a model system, prior work from our lab demonstrated that nucleotide sequence can play an independent role in driving functional distinctions of closely related protein isoforms [5]. The current study extends these observations to another cytoskeleton family and demonstrates that this principle also applies to tubulins isoforms. By extension, our findings imply that nucleotide- and amino acid-dependent determinants may constitute a common mechanism that underlies functional distinctions of multiple homologous isoform families and key essential proteins in vivo.

Our study shows that protein-level replacement of Nda2 with Atb2 leads to reduced microtubule dynamics, but does not result in strong phenotypic changes in the interphase cells or during mitosis – the processes that encompass the vegetative life cycle in yeast. In contrast, this tubulin replacement exerts dramatic effects on the meiotic spindle and meiotic chromosome segregation. While a number of important differences between mitosis and meiosis do exist [7], none of these previously identifies differences have been directly linked to tubulin, aside from the differences in the microtubule-organizing center. On the contrary, microtubule-dependent chromosome positioning and movements during mitosis and meiosis are believed to broadly follow similar steps. It is unclear what determinants could drive the differential functions of alpha tubulin during these events. Previous studies show that meiotic spindle, unlike the mitotic one, is acentrosomal, and it has been proposed that it assembles by kinesin-dependent “zipping” of microtubules to focus them at the poles, rather than by microtubule growth from the centrosomes [14, 15]. It is possible that this difference renders meiotic spindles more sensitive to perturbations, including changes in the balance of spindle motors and microtubule dynamics. In contrast, the mitotic spindles may be more robust and function normally under a broader range of conditions. Since sporulation and meiosis occurs in yeast much more rarely than mitosis, it likely exists under weaker evolutionary pressure and, by consequence, may be less robust compared to mitosis.

Previous studies outlined two major activities that play key roles in the spindle: the Ase1-dependent protein scaffolding that recruits many essential spindle components, and the microtubule end-binding proteins [10]. Our data show that both of these functions are perturbed in NCA yeast. The levels of Ase1 in the spindle midzone are reduced ∼2-fold, accompanied by the reduction of the Ase1-recruited kinesin, Klp9. The activity of the microtubule plus ends, tracked by Mal3, is also reduced. This suggests that the specific features of Nda2 amino acid sequence are uniquely required for the protein-protein interactions involved in the activity and function of these two protein complexes, and that substitutions of Nda2 sequence with that of Atb2 cannot fully support the same functions in cells.

Ase1 in the spindle acts in concert with other proteins, both via direct binding (such as Klp9) and indirect interplay that controls in the formation and maintenance of the metaphase bipolar spindle in meiosis (such as Klp2) [16]. To maintain their balance, both the absolute and the relative levels of these proteins in the spindle are precisely regulated. In the case of Klp9, this relationship is direct: reduced Ase1 in the spindle seen in NCA yeast is accompanied by a similar reduction of Klp9. This observation is in full agreement with the fact that Ase1 directly recruits Klp9. The situation with Klp2 is more complex, but potentially just as important. While Ase1 levels are reduced nearly 2-fold, the accompanying smaller change in Klp2 leads to its mild increase in the NCA spindles. Summarily, this change is expected to generates an imbalance between these two proteins in the spindle. Since Klp2 and Ase1 synergize to maintain meiotic spindle stability [16], reduced Ase1 and increased Klp2 binding capacity on meiotic spindle substantially alters their ratio. Conceivably, such an imbalance would likely lead to unstable metaphase spindle in NCA, and this instability could further cause the chromosome segregation defects observed in these cells. Klp9 is the major player for anaphase spindle elongation. Qualitative analysis of Klp9 showed roughly ∼2-fold reduction in its association with the meiotic spindle in NCA cells, this observation is consistent with the reduced anaphase velocity during meiosis.. At present, it is unclear how this meiosis-specific change could be regulated, but we speculate that this regulation could be driven by upstream regulatory factors – e.g., signaling molecules and protein modifying enzymes that may have different activity or target repertoire in meiotic versus mitotic cells. Since sporulation and meiosis occurs in yeast in response to major stimuli and environmental changes, it is highly likely that many signaling events could directly or indirectly target tubulin and its associated proteins by mechanisms that are not similarly employed in meiosis. Identification of molecular changes that differentially regulate mitosis and meiosis constitutes an exciting direction of further studies.

## MATERIALS AND METHODS

### Key resources table

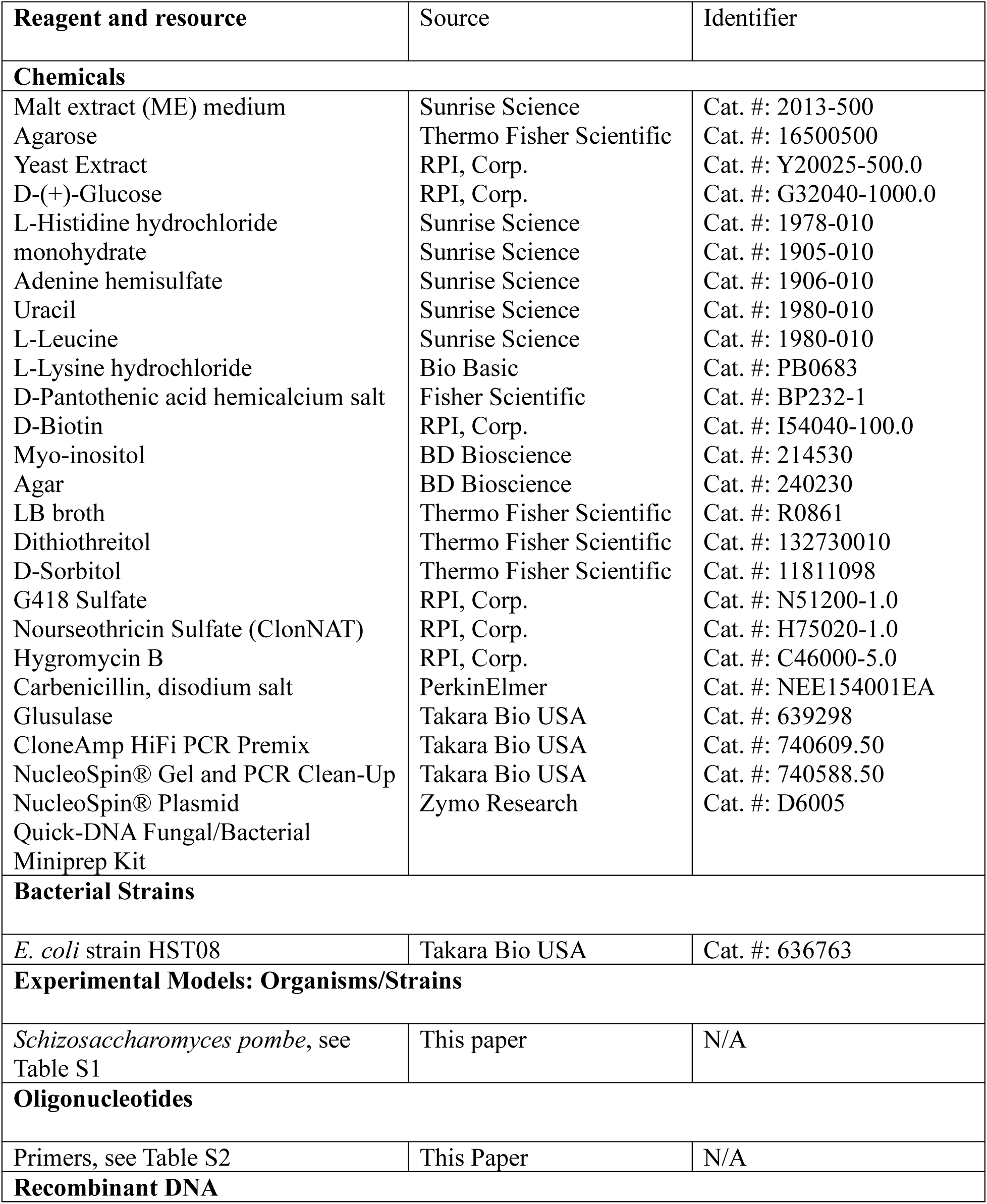
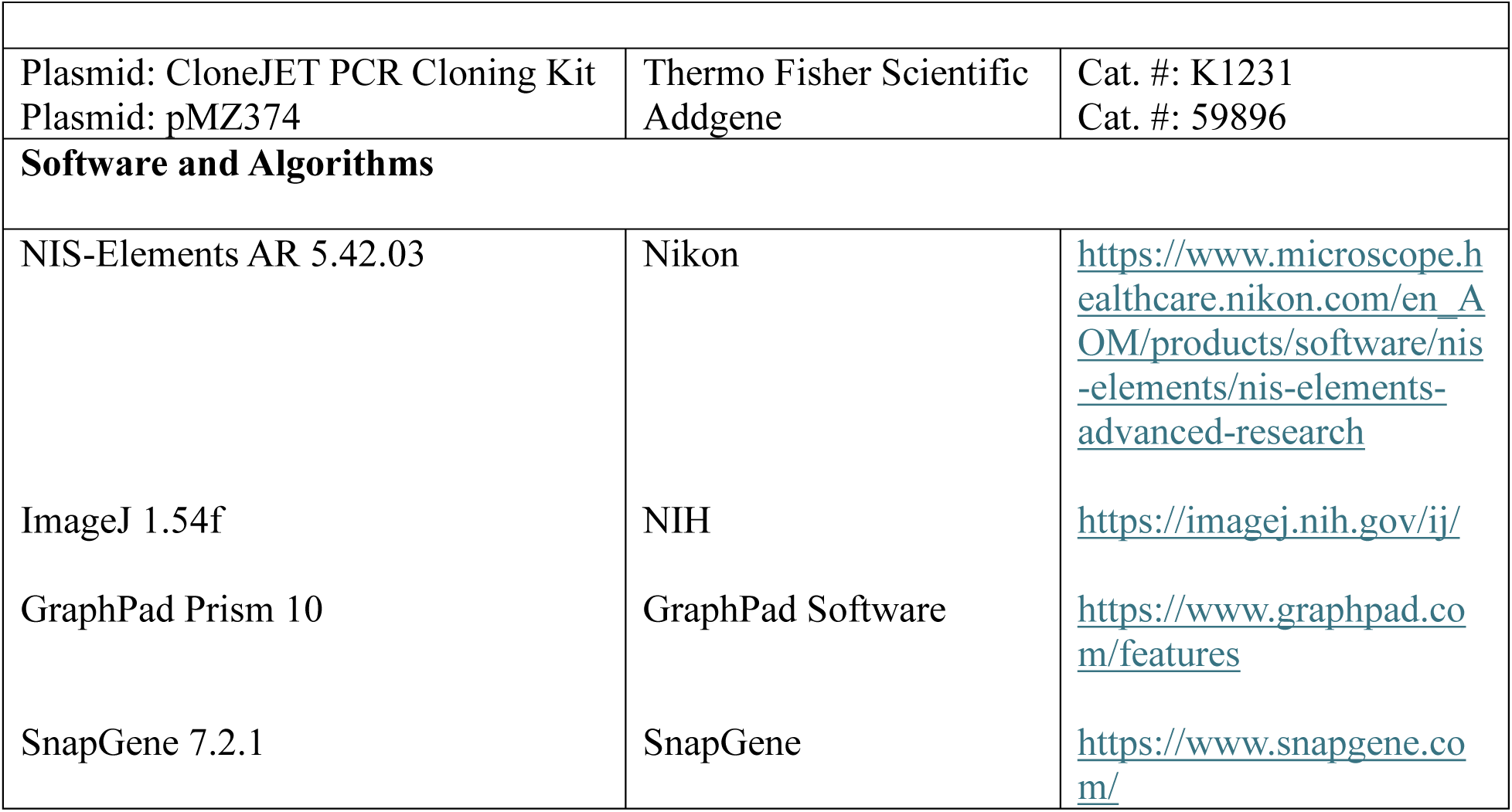

### Genetics, cell culture, and strains

Our study utilized the fission yeast *Schizosaccharomyces pombe* as the experimental model. A comprehensive list of all yeast strains used is provided in Table S1. Standard protocols for culture media and genetic manipulation were followed. Yeast cultures were typically grown at 25°C in YE5S medium (3% glucose, 0.5% yeast extract, 0.02% of Adenine, Uracil, Histidine, Leucine and Lysine, 2% agar for solid medium) [17]. SPA medium (1% glucose, 0.1% KH_2_PO_4_ 2% agar and 0.1% (v/v) Vitamins (1000×)) was used for yeast sporulation assay [17]. For live cell imaging analysis, YE5S (2% agarose instead of agar) and MEA medium (3% malt extract medium, 3% agarose) were used for vegetative and meiotic cells cultivation respectively during the whole imaging process. New strains were generated either by CRISPR-Cas mediated genome editing or by standard yeast genetics methods [18], [19]. Strains stably expressing proteins with GFP/mCherry tags were obtained by random spore germination followed by selection on plates with appropriate supplements or drugs.

### Plasmids

All primers were obtained from Integrated DNA Technologies, with details provided in Table S2. Constructs were sequenced at the DNA Sequencing Facility at the University of Pennsylvania. Plasmids of pMZ374 with sgRNAs targeting the up and down stream of nda2 untranslated regions (5’ and 3’ UTR) respectively were constructed according Jacobs, et al [18]. The donor template for *nda2*-coded *atb2* editing was inserted into the plasmid pJET 1.2 for co-transformation with pMZ374 containing sgRNAs. Electroporation mediated transformation was conducted in this research [20].

### Live-cell imaging and quantitative analysis

Images were acquired at room temperature (23°C) on a Nikon Eclipse Ti inverted microscope. The system, controlled by NIS-Elements AR software (Nikon), featured a Yokogawa CSUX1 spinning-disk confocal head, a Nikon 100×/1.49 NA Plan Apochromat oil immersion lens, and an Andor iXon Ultra 897 EMCCD camera. For vegetative cells analyses, the cells were cultured overnight until they reached an optical density at 600 nm (OD_600_) between 0.4 and 0.8.

Afterward, the cells were harvested and placed on 2% agarose pads made with YE5S medium for live-cell imaging. Spindle dynamics graphs were obtained by analysis of spindle pole distance over time, nuclear envelope breakdown indicated the end of spindle elongation (images were taken at 1-min intervals with 7 optical sections of 1-*μ*m spacing). For meiosis analyses, the two different mating type cells h+ and h-were cultivated overnight respectively until they reached an optical density at 600 nm (OD_600_) between 0.4 and 0.8. Then mixed same amount of each type of cells followed by centrifugation to discard the supernatant, the pellet was washed twice by ME broth medium. The mixed pellet of opposite mating type cells was cultivated on SPA medium for ∼18 h, then the cells were collected and placed on 3% agarose pads made with ME medium for live-cell imaging (images for spindle dynamics were taken at 2-min intervals, for chromosome segregation assay were taken at 3-min intervals. All images were captured with 7 optical sections of 1-*μ*m spacing). Meiotic interphase microtubule (MT) dynamics were captured at 5-second intervals across three optical sections with 1-μm spacing. The rates of MT growth were calculated by tracking changes in MT binding protein Mal3-GFP movement from time-lapse sequences. The dwell time was defined as the duration during which the Mal3-GFP signal remained in contact with the cell tip before disappearing. Fluorescence intensities were measured as total intensity, with background subtracted, within a defined region that represented the sum projection of all focal planes encompassing the entire cell.

All graphs were created using GraphPad Prism 10.0. Column graphs with scatter plot display mean values with error bars indicating ±SD. Statistical significance was assessed using unpaired T tests.

## ACKNOWLEDGEMENTS

We thank Prof. Phong Tran for helpful discussions and the gift of yeast strains and plasmids with GFP protein markers, and members of the Kashina lab, especially Dr. Junling Wang, Dr. Pavan Vedula, and Brittany MacTaggart for their input and feedback throughout the project. This work was supported by NIH grants R35GM122505 and R01NS102435 to AK.

## AUTHOR CONTRIBUTIONS

L.C. designed and performed experiments, analyzed data, wrote the paper; X.C. performed experiments; A.K. designed experiments, analyzed data, wrote the paper.

## SUPPLEMENTAL ONLINE INFORMATION

**Table S1.** Fission yeast strains used in this study.

| Strains | Genotype | Source |
| --- | --- | --- |
| PT.286 | h- / WT | Phong Tran |
| PT.287 | h+ / WT | Phong Tran |
| CL.107 | h- / Nda2 coded Atb2 (NCA with intron) | This paper |
| CL.108 | h- / Nda2 coded Atb2 (NCA with intron) | This paper |
| CL.110 | h+ / Nda2 coded Atb2 (NCA with intron) | This paper |
| CL.111 | h+ / Nda2 coded Atb2 (NCA with intron) | This paper |
| CL.127 | h- / Sid4-mCherry:HygR / Cut11-GFP:URA4 | This paper |
| CL.128 | h+ / Sid4-mCherry:HygR / Cut11-GFP:URA4 | This paper |
| CL.133 | h+ / NCA with intron / Sid4-mCherry:HygR / Cut11-GFP:URA4 | This paper |
| CL.134 | h- / NCA with intron / Sid4-mCherry:HygR / Cut11-GFP: URA4 | This paper |
| CL.135 | h- / NCA with intron / Sid4-mCherry:HygR / Cut11-GFP: URA4 | This paper |
| CL.136 | h+ / NCA with intron / Sid4-mCherry:HygR / Cut11-GFP: URA4 | This paper |
| CL.115 | h+ / Hht2-GFP:URA4 | Phong Tran |
| CL.129 | h- / Hht2-GFP:URA4 | This paper |
| CL.123 | h+ / NCA with intron / Hht2-GFP:URA4 | This paper |
| CL.124 | h- / NCA with intron / Hht2-GFP:URA4 | This paper |
| CL.125 | h+ / NCA with intron / Hht2-GFP:URA4 | This paper |
| CL.126 | h- / NCA with intron / Hht2-GFP:URA4 | This paper |
| CL.173 | h+ / NCA with intron / Sid4-mCherry:HygR / Klp9-GFP:NatR | This paper |
| CL.174 | h- / NCA with intron / Sid4-mCherry:HygR / Klp9-GFP:NatR | This paper |
| CL.175 | h+ / NCA with intron / Sid4-mCherry:HygR / Klp9-GFP:NatR | This paper |
| CL.176 | h- / NCA with intron / Sid4-mCherry:HygR / Klp9-GFP:NatR | This paper |
| CL.181 | h+ / Sid4-mCherry:HygR / Klp9-GFP:NatR | This paper |
| CL.182 | h- / Sid4-mCherry:HygR / Klp9-GFP:NatR | This paper |
| CL.203 | h+ / NCA with intron / Sid4-mCherry:HygR / Klp2-GFP:URA4 | This paper |
| CL.204 | h- / NCA with intron / Sid4-mCherry:HygR / Klp2-GFP:URA4 | This paper |
| CL.205a | h+ / NCA with intron / Sid4-mCherry:HygR / Klp2-GFP:URA4 | This paper |
| CL.205b | h- / NCA with intron / Sid4-mCherry:HygR / Klp2-GFP:URA4 | This paper |
| CL.206 | h- / NCA with intron / Sid4-mCherry:HygR / Ase1-GFP:URA4 | This paper |
| CL.207 | h+ / NCA with intron / Sid4-mCherry:HygR / Ase1-GFP:URA4 | This paper |
| CL.208 | h- / NCA with intron / Sid4-mCherry:HygR / Ase1-GFP:URA4 | This paper |
| CL.209 | h+ / NCA with intron / Sid4-mCherry:HygR / Ase1-GFP:URA4 | This paper |
| CL.220 | h- / Sid4-mCherry:HygR / Klp2-GFP:URA4 | This paper |
| CL.221 | h+ / Sid4-mCherry:HygR / Klp2-GFP:URA4 | This paper |
| CL.222 | h+ / Sid4-mCherry:HygR / Ase1-GFP:URA4 | This paper |
| CL.223 | h- / Sid4-mCherry:HygR / Ase1-GFP:URA4 | This paper |

**Table S2.**
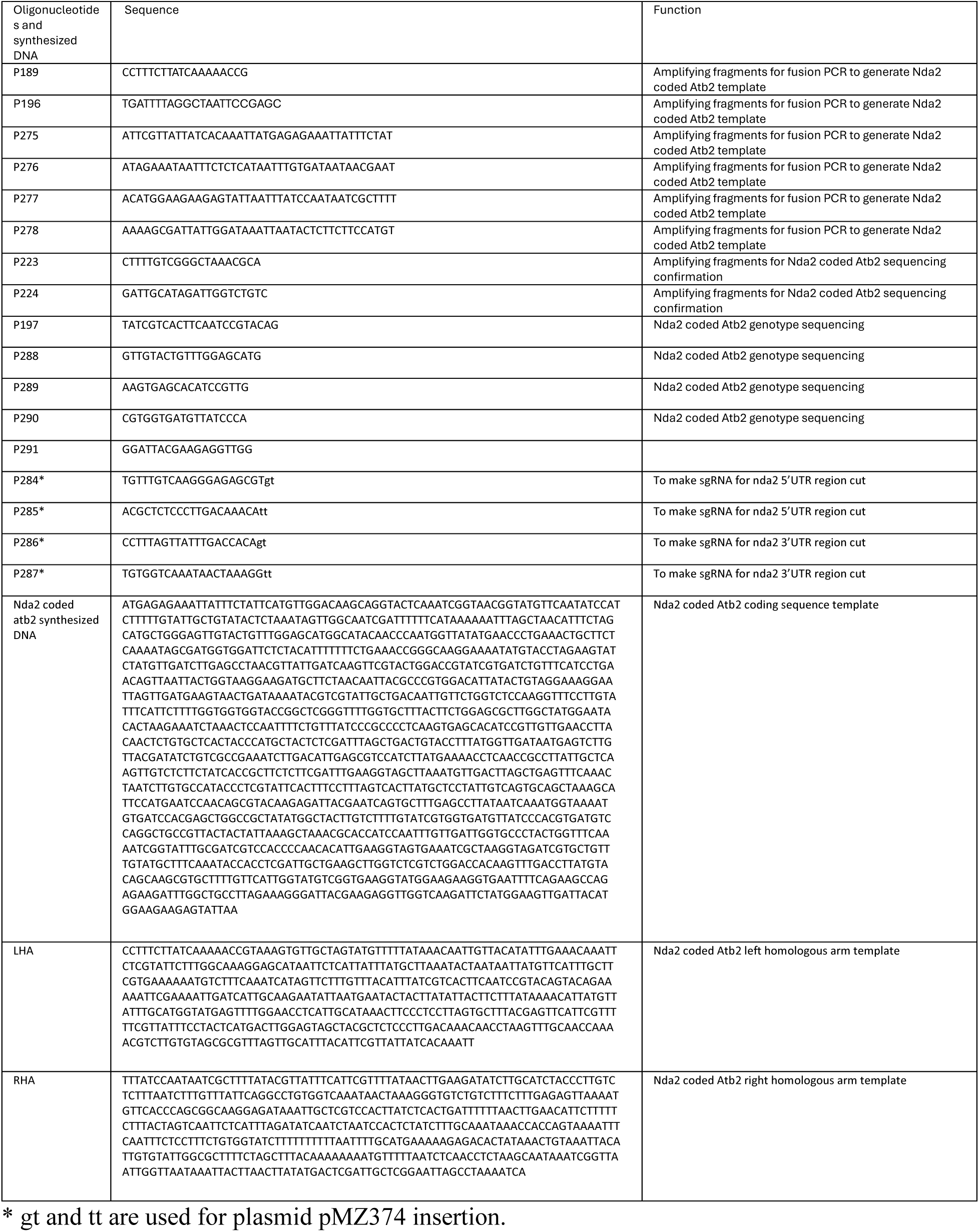
Oligonucleotides and synthesized DNA used in this study.

**Figure S1.**
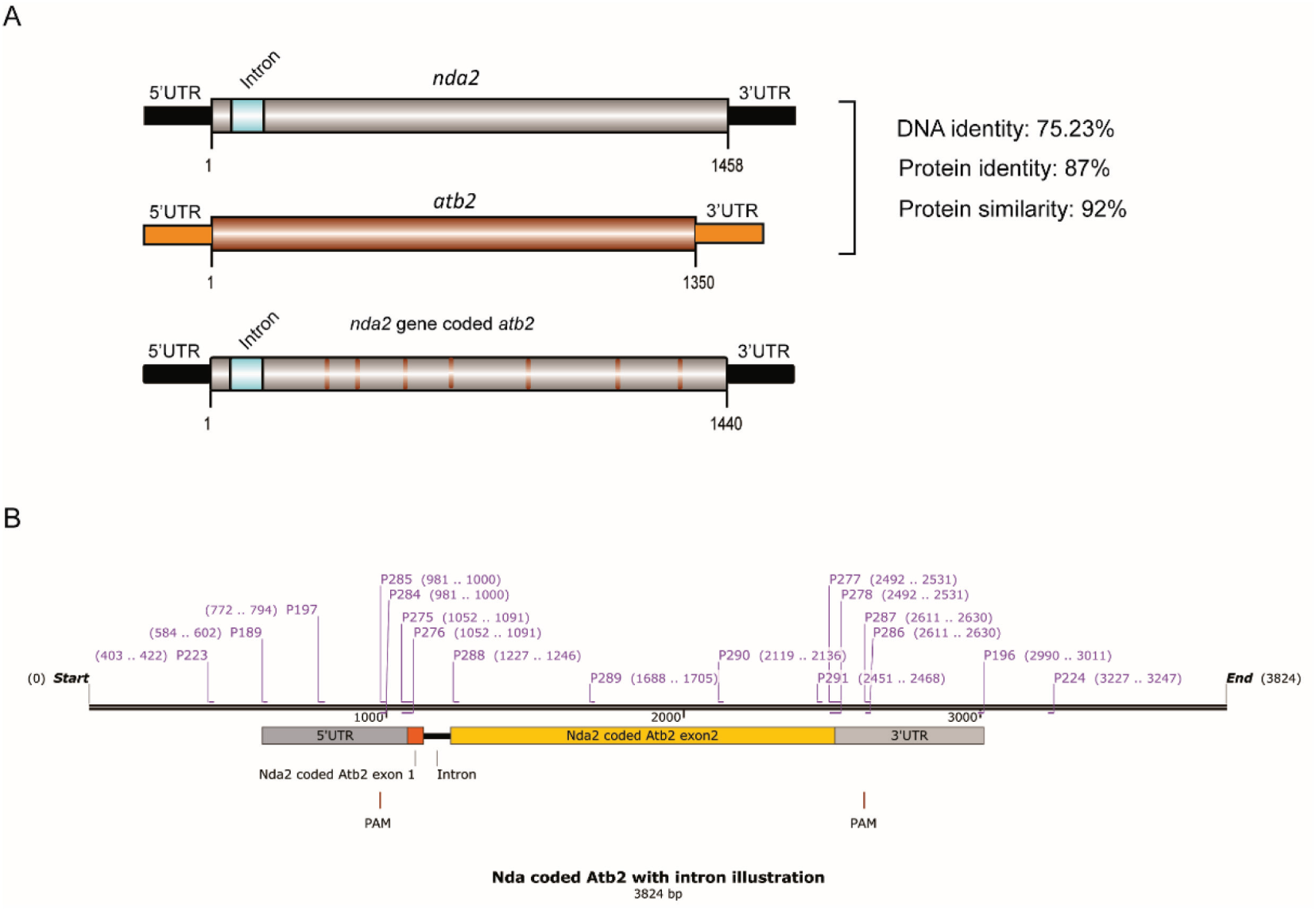
Illustration of Nda2, Atb2 and Nda2 coded Atb2. (A) Comparison of Nda2 and Atb2 on nucleic acid and protein level. (B) Illustration of gene structure of Nda2 coded Atb2.

